# The structured diversity of specialized gut symbionts of the New World army ants

**DOI:** 10.1101/084376

**Authors:** Piotr Łukasik, Justin A. Newton, Jon G. Sanders, Yi Hu, Corrie S. Moreau, Daniel J. C. Kronauer, Sean O’Donnell, Ryuichi Koga, Jacob A. Russell

## Abstract

Symbiotic bacteria play important roles in the biology of their arthropod hosts. Yet the microbiota of many diverse and influential groups remain understudied, resulting in a paucity of information on the fidelities and histories of these associations. Motivated by prior findings from a smaller scale, 16S rRNA-based study, we conducted a broad phylogenetic and geographical survey of microbial communities in the ecologically dominant New World army ants (Formicidae: Dorylinae). Amplicon sequencing of the 16S rRNA gene across 28 species spanning the five New World genera showed that the microbial communities of army ants consist of very few common and abundant bacterial species. The two most abundant microbes, referred to as Unclassified Firmicutes and Unclassified Entomoplasmatales, appear to be specialized army ant associates that dominate microbial communities in the gut lumen of three host genera, *Eciton, Labidus* and *Nomamyrmex*. Both are present in other army ant genera, including those from the Old World, suggesting that army ant symbioses date back to the Cretaceous. Extensive sequencing of bacterial protein-coding genes revealed multiple strains of these symbionts co-existing within colonies, but seldom within the same individual ant. Bacterial strains formed multiple host species-specific lineages on phylogenies, which often grouped strains from distant geographic locations. These patterns deviate from those seen in other social insects, and raise intriguing questions about the influence of army ant colony swarm-founding and within-colony genetic diversity on strain co-existence, and the effects of hosting a diverse suite of symbiont strains on colony ecology.

## Introduction

Insects are the world’s most speciose group of animals and have come to dominate most terrestrial ecosystems. Growing evidence suggests that this dominance has, in many cases, been facilitated by their microbial associates, which can expand their hosts’ metabolic repertoires and ranges of environmental tolerance. Symbionts of insects represent a broad diversity of bacterial and fungal lineages, and vary tremendously in their distributions within host bodies, their biological effects, their transmission modes and their co-evolutionary histories with hosts (Douglas 2015; McFall-Ngai *et al.* 2013; Moran *et al.* 2008).

The often dramatic effects of insect endosymbionts – bacteria that colonize host tissues and cells – are particularly well understood. Obligate nutritional endosymbionts have enabled major nutritional transitions onto imbalanced diets, resulting in highly speciose lineages such as Hemiptera (Baumann 2005; Bennett & Moran 2015). Facultative endosymbiotic bacteria can also have dramatic effects on life history traits of hosts, including their reproductive biology or susceptibility to natural enemies (Feldhaar 2011; Oliver *et al.* 2010). This can cause major changes to insect populations (Himler *et al.* 2011; Jaenike *et al.* 2010) that can resonate across communities (Ferrari & Vavre 2011; Sanders *et al.* 2016). However, insects commonly host bacteria outside of their haemocoel – on their cuticles or within the digestive tract (Engel & Moran 2013). The size, composition, location and functions of these ectosymbiotic microbial communities vary considerably across taxa (Engel & Moran 2013). In some cases, bacterial numbers are relatively low, and microbial communities consist of genera commonly found in the environment (Broderick *et al.* 2014; Chandler *et al.* 2011). However, in many hosts ectosymbiont communities are remarkably stable across individuals, populations and over longer evolutionary timescales, with the key members showing strong host fidelity and, possibly, only rare host switching (Hu *et al.* 2014; Koch *et al.* 2013; Kwong & Moran 2015; Sanders *et al.* 2014; Sudakaran *et al.* 2012). In at least some cases, these microbes colonize only selected host tissues (Kautz *et al.* 2013b; Lanan *et al.* 2016; Martinson *et al.* 2012) and the microbial communities change in predictable ways as their hosts develop (Sudakaran *et al.* 2012). Knowledge on the functions of ectosymbionts still lags behind that for endosymbionts. However, recent advances in the field of genomics, as well as experimental approaches, have revealed diverse effects on numerous aspects of host biology, including digestion, detoxification, nutrition, susceptibility to parasites and pathogens, development, regulation of host physiology and communication with members of the same or other species (Engel & Moran 2013).

As seen for endosymbionts (Duron *et al.* 2008; Russell *et al.* 2012), early studies of ectosymbionts often involved diagnostic PCR screening or Sanger-sequencing-based cloning of small numbers of sequences to characterize microbiota (e.g., Corby-Harris *et al.* 2007; Russell *et al.* 2009b). Other efforts focused on cultured representatives from ectosymbiont communities, raising questions about the impacts of culture bias on our understanding of these symbioses (e.g., Broderick *et al.* 2004). The adoption of 16S rRNA gene amplicon sequencing on next-generation platforms has greatly expanded the depth of culture-independent community sampling (Jones *et al.* 2013; Yun *et al.* 2014). However, the low rate of evolution of this gene limits our abilities to detect cryptic, strain-level variation (Ellegaard & Engel 2016). Indeed closely related, conspecific bacteria exist in numerous ectosymbiont communities, including those found in the guts of bees, ants and termites (Ellegaard & Engel 2016; Engel *et al.* 2012; Engel *et al.* 2014; Hu *et al.* 2014; Powell *et al.* 2016; Warnecke *et al.* 2007). And as seen for bacterial endosymbionts of insects (e.g., Oliver *et al.* 2005), the functions of closely related ectosymbiont strains can differ (Engel *et al.* 2012). To truly understand ectosymbiont diversity and the relationship between the microbiota and host biology, researchers should target conserved genes with a history of vertical transfer (i.e. 16S rRNA) to aid in symbiont classification, but also more variable loci to differentiate strains.

The diversity and function of ectosymbiont communities have garnered a good deal of recent attention among the ants (Hymenoptera: Formicidae), a diverse and ecologically dominant group of social insects that colonize a wide range of trophic and habitat niches (Davidson *et al.* 2003; Holldobler & Wilson 1990). Specialized symbionts have been implicated as important partners of ants with specialized diets. For example, ectosymbiotic bacteria function as biocontrol agents in the management of attine ants’ fungal gardens (Currie *et al.* 1999), and recent studies have placed an emphasis on the diversity and structure of these communities (Andersen *et al.* 2013).

While at least one group of herbivorous ants has evolved a monotypic nutritional symbiosis with intracellular bacteria (Feldhaar *et al.* 2007; Wernegreen *et al.* 2009), others harbor substantial ectosymbiont populations in their gut cavities (Billen & Buschinger 2000; Roche & Wheeler 1997; Sanders *et al.* in review). Hypothesized to have shaped the ants’ adaptation to plant-based diets (Davidson et al. 2003; Russell et al. 2009b), ectosymbiotic gut bacteria generally vary across herbivorous ant taxa (Anderson *et al.* 2012; Sanders *et al.* 2014; Stoll *et al.* 2007), exhibiting modest diversity at the species and strain levels (Hu et al. 2014). As seen for the fungus-growing ants (Andersen *et al.* 2013), strain-level composition of these communities appears similar across sibling workers yet variable between colonies (Hu et al. 2014).

It has not yet been established whether the stereotyped and stable ectosymbiont communities of fungus-growers (cuticular bacteria) and liquid-feeding ant herbivores (gut bacteria) represent the norm across this insect family. In fact, some ants appear to harbor few bacteria while others show much greater variability in their ectosymbiotic compositions (Liberti *et al.* 2015; Russell *et al.* accepted; Sanders *et al.* in review). It is notable, then, that predatory army ants represent a third guild of trophic specialists engaging in specialized, multi-partite ectosymbioses (Anderson *et al.* 2012; Funaro *et al.* 2011). Army ants are keystone species in many of the world’s tropical forests (Boswell *et al.* 1998; Kaspari & O’Donnell 2003), living in large colonies with one queen and anywhere from tens of thousands to millions of workers (Kronauer 2009). Unlike fungus-growers and herbivorous ants such as *Cephalotes* and *Tetraponera*, new army ant colonies are always founded by colony fission, where swarms made up by a single, mated queen and tens to hundreds of thousands of accompanying workers depart from the maternal colony (Cronin *et al.* 2013; Kronauer 2009; Peeters & Ito 2001). These differences in modes of colony founding across known hosts of specialized symbionts raise questions about the diversity of microbial communities, but also how such diversity is partitioned among versus within colonies.

The goal of this study was to characterize the composition and structure of bacterial communities of the New World Dorylinae in a comprehensive, multi-level fashion, across individuals, colonies, species, as well as geography. To achieve this, we used multiplex Illumina sequencing of 16S rRNA amplicons for a total of 195 individual workers from 102 New World army ant colonies representing 28 species from the five described genera. Amplicon sequencing of 16S rRNA was also performed for colonies representing two doryline army ant genera from the Old World. For the two most broadly sampled New World species, *Eciton burchellii* and *Labidus praedator*, we targeted multiple workers from replicate colonies collected at different geographic locations. We then characterized strain-level diversity and host specificity of the two dominant symbiont species by sequencing the ribosomal protein L2 (*rplB*) gene for approximately 350 bacterial strains. Finally, we used fluorescent *in-situ* hybridization to visualize specific symbionts within army ant tissues, yielding the first clear insights into their localization.

## Materials and Methods

### Ant collections

The 102 Dorylinae colonies used in the study were collected between 2001 and 2013 in six countries in the Americas: Costa Rica, Ecuador, Mexico, Peru, Venezuela and U.S.A., typically from more than one locality within each country (Table S1). Two sites, Henri Pittier National Park (Venezuela) and the Monteverde area (Costa Rica), were the most extensively sampled. The collected ants were identified to the genus or species level based on morphology and colony characteristics, and then preserved in ethanol or acetone and stored at −20°C until further processing.

We also investigated microbial communities of specimens from 15 Old World army ant colonies used by Funaro et al. (2011; Table S2). In some cases, intact ethanol-preserved workers were available, and they were processed the same way as workers from the New World. In other cases we used DNA extracted following different protocols by Funaro et al. (2011).

### DNA extraction

From each colony, we aimed to process five medium-sized individual workers whenever possible (Table S3). Insects were surface-sterilized by immersion in 5% bleach for 60 seconds and then washed three times with molecular-grade water. Next, from each worker we dissected the gaster, the posterior part of the body that contains all sections of the digestive system other than esophagus. We took care to remove any fragments of the petiole (i.e. the body section immediately anterior to the gaster), and to sterilize forceps between dissections. Individual gasters were flash-frozen in liquid nitrogen, ground with a pestle, and then processed using DNeasy Blood and Tissue kits (Qiagen Ltd., Hilden, Germany) following the manufacturer’s protocol for Gram-positive bacteria. In order to control the level of background contamination with bacterial DNA, we included ‘blank’ samples in all extraction batches, typically one for every ten ants processed.

### Molecular identification of ants

For one worker from each colony we attempted to amplify and sequence a part of the cytochrome oxidase I (*COI*) gene. We first attempted to amplify a ~1100bp fragment using universal primers LCO-1490 and BEN, and sequence it with BEN and internal primer HCO-2198 (Table S4). For samples for which we failed to generate product or for which sequence quality was low, we used primers LCO-1490 and HCO-2198 to amplify and sequence ~650bp of the gene. A range of PCR conditions were used (Table S4). However, for 15 colonies our attempts to generate unambiguous *COI* sequences were unsuccessful. All traces were manually inspected, assembled, edited and aligned using CodonCode Aligner v. 4.2.7 (CodonCode Corp., Centerville, MA).

### Microbial community analysis

All extracted samples, including ‘extraction blanks,’ were used for PCRs with universal eubacterial primers for the 16S rRNA gene, 9Fa and 907R, conducted under standardized conditions (Table S4). Concentrations of bacterial DNA in the samples were roughly estimated by running 5 µl of the PCR product on a 2% agarose gel stained with ethidium bromide and comparing the brightness of the bands against the negative controls. Samples yielding strong *COI* bands but with 16S rRNA bands of similar or lesser strength to those for our negative controls were re-tested with other pairs of eubacterial primers. When such primers failed to amplify, these samples were classified as hosting insufficient numbers of bacteria for reliable microbial community characterization (Table S3), and were not analyzed further.

The composition of microbial communities of ants hosting substantial numbers of bacteria was characterized using amplicon sequencing of the V4 hypervariable region of the 16S rRNA gene on an Illumina MiSeq platform, conducted following the Earth Microbiome Project protocols (Caporaso et al. 2012). Library preparation and sequencing were conducted at Argonne National Laboratory (Argonne, IL). For New World Dorylinae, we submitted DNA samples from a single worker gaster from each colony, whenever there was at least one exceeding the rough bacterial density threshold specified above. Furthermore, to investigate bacterial distributions at both the colony and species levels, we selected four additional workers from each of 12 colonies of *Labidus praedator* and from each of 15 colonies of *Eciton burchellii*. Our submission also included 15 samples from the Old World army ants (genera *Aenictus* and *Dorylus*). Finally, we included 13 blanks: either ‘extraction blanks’, or molecular grade water. Data were analyzed using Mothur v 1.36.1 (Schloss et al. 2009). Analysis steps are summarized below, details and specific commands are provided in the Supplementary Material.

After merging quality-filtered and trimmed reads into contigs and then identifying unique genotypes, we used a custom Python script to conduct a custom decontamination procedure, based on the abundance of unique genotypes in experimental and blank libraries. For each unique 16S rRNA genotype identified through sequencing, we compared the maximum relative abundance it attained in any of the blanks and in any of the experimental libraries; we classified sequences as contaminants when their unique genotypes were not at least ten times more abundant in at least one experimental library compared to the maximum abundance seen across the blanks. Subsequently, we removed any ‘rare’ unique genotypes, which did not constitute at least 0.1% of at least one experimental library. Finally, those experimental libraries that lost more than 75% of reads during the previous steps were deemed heavily contaminated and also removed. The decontamination procedure is described in more detail in the Supplementary Material, and its effects on the microbial community composition are summarized in Figure S1. Reads were then aligned against a custom set of 16S rRNA references, screened for chimaeras with UChime, and classified using the RDP training set (v. 9). All reads that did not classify as Eubacteria were subsequently removed.

The resulting dataset was used for OTU clustering at the 97% identity level using the average neighbor algorithm. We then manually processed the output files (i.e. the alignment of all unique sequences, information about taxonomic classification of each, the OTUs to which each unique sequence was assigned, and the number of times each unique sequence appeared in each of the libraries) – using Microsoft Excel and custom Python and Processing scripts. We used ADONIS, from the vegan package (Oksanen *et al.* 2007) in R v. 3.1.3 (R Core Team 2015), to compare the composition of the microbial communities between the two most extensively sampled species, *Eciton burchellii* and *Labidus praedator*, across collection locations, and among colonies of the same species within locations. In case of significant differences, we implemented ANOVA tests in QIIME v. 1.9.1 (Caporaso *et al.* 2010) to identify OTUs that differed in relative abundance among groups in a comparison.

### Sequence typing of the dominant symbionts

We investigated the phylogenetic relationships of the two dominant symbionts of army ants, Unclassified Firmicutes and Unclassified Entomoplasmatales. The relationship of Unclassified Firmicutes with other bacterial taxa was assessed using 16S rRNA gene sequences, for which more extensive comparative data were available. Nearly full-length 16S rRNA sequences of Unclassified Firmicutes were obtained by direct sequencing of PCR products obtained with universal eubacterial primers 9Fa and 1513R for ants dominated by a single bacterial strain, or with diagnostic primers UNF-16S-F1 and UNF-16S-R1 for all others (Table S4). Using these protocols we amplified 16S rRNA gene from strains infecting 19 ants, including Old World army ants and the two Ponerinae species previously identified to host Unclassified Firmicutes (Tables S2–S3). PCR products were purified by digestion with Exonuclease I and Antarctic Phosphatase (NEB, Ipswich, MA) and sequenced by Eurofins MWG Operon (Louisville, KY). To characterize the relationships between Unclassified Firmicutes strains, we developed a protocol for amplifying and sequencing the terminal 579bp fragment of a conserved single-copy gene encoding ribosomal protein L2 (*rplB*; Table S4). The reads were manually inspected, quality-trimmed and aligned using CodonCode Aligner. We successfully sequenced *rplB* from 252 Unclassified Firmicutes strains, with representation from all Dorylinae genera. Special emphasis was placed on the two most broadly sampled ant species, *Eciton burchellii* and *Labidus praedator*, for which numerous sequences were generated per colony and site (Tables S1, S3).

Phylogenetic relationships of army ant-associated Entomoplasmatales had been inferred previously using the slowly evolving 16S rRNA gene (Funaro *et al.* 2011). To gain further insight into the strain-level diversity within this group, we developed a protocol for amplifying a 573bp segment of *rplB* from Entomoplasmatales (Table S4), using the generated amplicons for Sanger sequencing across 120 army ants and a small number of other ants and insects (Table S2).

### Genotype-level diversity of Unclassified Firmicutes in individual workers

We used 454 pyrosequencing of *rplB* amplicons to assess strain-level diversity of Unclassified Firmicutes in four workers from *Eciton burchellii* colony PL028. Sanger sequencing results had suggested that each of the selected workers was dominated by a different symbiont strain, or genotype, spurring us to perform deep sequencing to more thoroughly assess within-host strain diversity. Products for pyrosequencing were generated using the same *rplB* primers as for PCRs prior to Sanger sequencing, complete with linkers and barcodes. These products were then sequenced in one direction on a multiplexed lane by Research and Testing Laboratory (Lubbock, TX). The data were analysed using Mothur v. 1.33 (Schloss *et al.* 2009). Quality-trimmed reads were aligned against Sanger sequences for the dominant strains in the four samples. We then clustered reads into OTUs at the 98% sequence identity level. This threshold was chosen as the lowest sequence similarity cut-off allowing us to distinguish between the dominant Unclassified Firmicutes genotypes in the four studied workers (in one of the pair-wise comparisons, genotypes had 97.5% identity across 360 bases). Further analysis details are provided in the Supplementary Material.

### Phylogenetic analyses

Sequences were unambiguously aligned using ClustalW in CodonCode Aligner. Maximum likelihood phylogenies of army ant mtDNA (i.e. partial *COI* sequences) and genes from their two dominant symbionts (*rplB* for both, and 16S rRNA for Unclassified Firmicutes) were reconstructed using RAxML 8.2.0 (Stamatakis 2014). In all cases we specified the GTR model with the addition of invariant sites and gamma distribution of rates across sites, one of the top three sequence evolution models for each of these datasets according to jmodeltest2 (Darriba *et al.* 2012). One thousand bootstrap replicates were run for each search. Host and symbiont phylogenies were illustrated using iTOL v.3.2.1 (Letunic & Bork 2016).

### Fluorescence in situ hybridization

We investigated the localization of symbiotic bacteria within the digestive tract of ants using fluorescence microscopy. Guts dissected from 6-10 workers from colonies of *Eciton rapax* (PL150), *Nomamyrmex esenbeckii* (PL157) and *Labidus praedator* (PL158) were fixed in 4% formaldehyde. Then, they were either used for whole-mount hybridization, or embedded in glycol methacrylate resin (Technovit 7100 - Heraeus Kulzer GmbH, Wehrheim, Germany) and sectioned prior to hybridization. Additionally, digestive tracts dissected from workers of *Eciton burchellii* colony JSC093 were used for whole-mount FISH only. We also studied a small number of specimens from six other colonies of four Dorylinae species, but because of their initial preservation in ethanol or acetone, tissue structure was not well preserved. Seven ant species from other subfamilies were processed using the same methods, thus serving as controls (Sanders *et al.* in review).

We used a combination of eubacterial probes (EUB338 and EUB897), as well as probes specific to Unclassified Firmicutes and Entomoplasmatales. The specific probes were originally developed as diagnostic and sequencing primers for the two specialized army ant symbionts, and tested extensively in a range of PCR conditions. The specificity of fluorescent signal was demonstrated in control experiments, where hybridization was done with no probes, or probes labeled with the same fluorophores but for bacteria that were believed to be absent in the sample. The signal of both probes was visibly increased when they were used in combination with helper probes – unlabeled oligos for the rRNA region immediately adjacent to that targeted by the probe (Fuchs *et al.* 2000). Finally, for the Entomoplasmatales probe, we verified that its signal overlapped with the signal of two other, previously verified probes for a bacterium from that order, *Spiroplasma* (TKSspi and TKSSsp - Fukatsu & Nikoh 2000), which were a perfect or near-perfect match to our sequences. All probe sequences are provided in Table S5. The detailed FISH protocol is provided as a supplement.

## Results

### Ant species identification and phylogenetics

We obtained reliable *COI* sequence data for workers from 87 out of 102 experimental New World Dorylinae colonies. Some of the colonies for which we failed to obtain unambiguous sequence data were confidently classified based on morphology to the same species as colonies with *COI* data from the same site (Table S1). However, reliable data could not be obtained for four species-location combinations. Their approximate position on the *COI* tree was indicated using dashed lines (Fig. 1).

**Figure 1.**
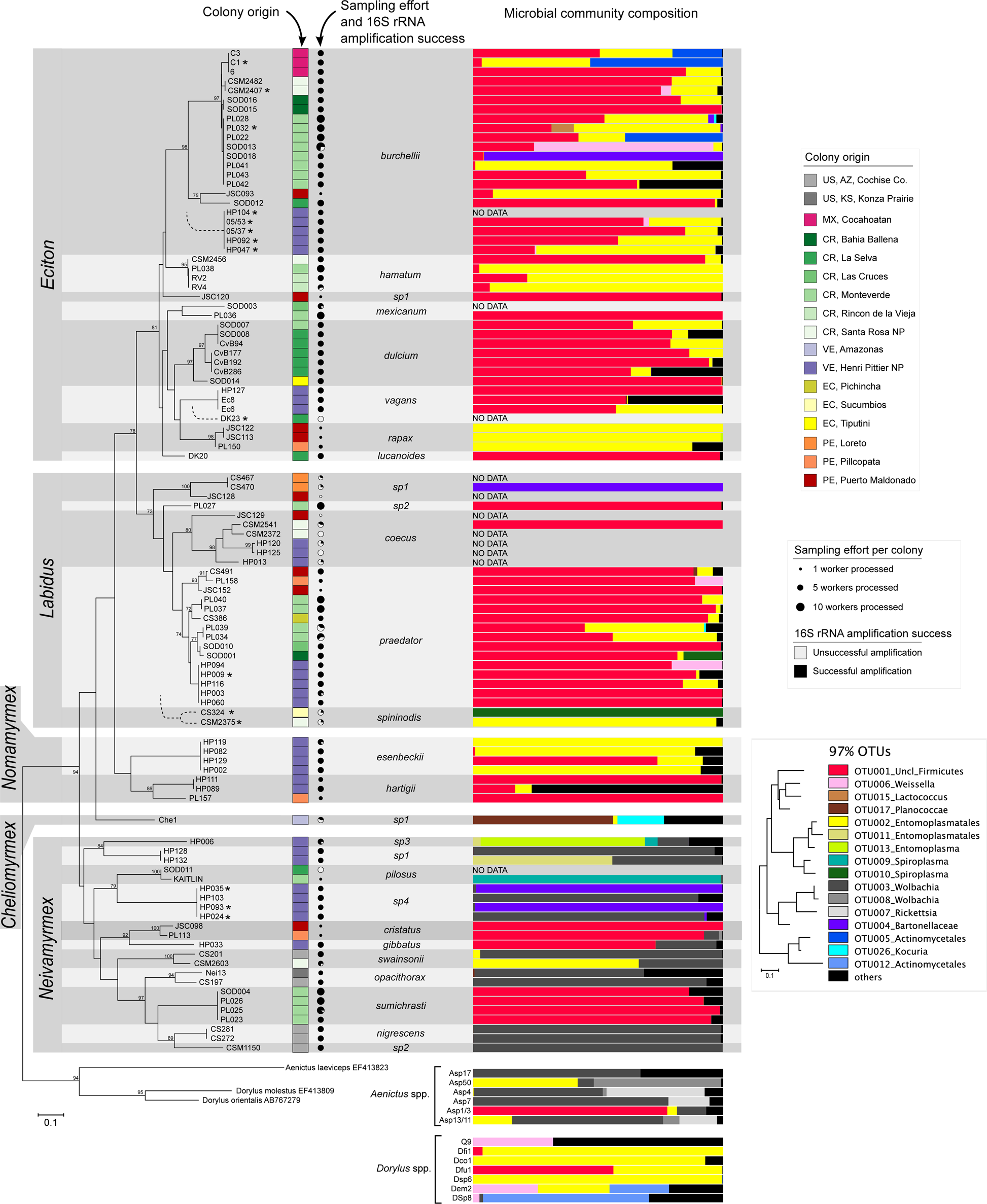
The diversity of microbial symbionts across New World army ants. The ML mtDNA phylogeny of Dorylinae (all genotype/location combinations in our collection) is based on 600bp sequence of the mitochondrial *COI* gene, and constrained so that *Neivamyrmex* spp. form a clade (following Brady 2003). Branch lengths represent the mean number of nucleotide substitutions per site. Bootstrap support values of 70% or more are shown as numbers above nodes. Colonies for which no unambiguous sequence data were obtained (indicated with asterisks) were assumed to be identical at *COI* to morphologically identical colonies from the same location. For four species-location combinations with no available sequence data, their approximate position on the tree is indicated with dashed lines. For all colonies, we indicate their geographic origin, the number of individual workers with DNA extracted, and the proportion of workers for which we succeeded in amplifying 16S rRNA. The relative abundance of bacterial 97% OTUs in ant gasters, based on 16S rRNA amplicon sequencing, is shown for the one worker characterized from each colony, except for those colonies of *E. burchellii* and *L. praedator* which are featured in Figure 2, where we used an average for all characterized siblings from a single colony. The relationship between symbiont OTUs is demonstrated using a ML phylogeny for the most abundant genotypes from each OTU.

The reconstructed *COI* tree provided a good overview of the sampled ant diversity (Fig. 1), recapitulating several previously documented relationships recovered with larger sequence datasets. At the same time, several deeper nodes of the tree disagreed with these prior studies, due likely to the insufficiency of a single genetic marker in accurate phylogenetic reconstruction. More specifically, samples classified to the same species based on morphology formed monophyletic clades, typically with high bootstrap support. The reconstructed relationships between species within the genus *Eciton* did not conflict with the results of a recent phylogenomic study (Winston *et al.* 2016). However, according to Brady (2003) and Borowiec (2016), the genus *Neivamyrmex* is monophyletic, and thus we constrained the *COI* tree to enforce *Neivamyrmex* monophyly. Furthermore, recent phylogenetic reconstructions based on multiple genes or genomic datasets (Borowiec 2016; Brady *et al.* 2014; Winston *et al.* 2016) identified *Nomamyrmex* as the sister genus of *Eciton*, whereas in the *COI* tree the genera *Eciton* and *Labidus* formed a monophyletic clade, with *Nomamyrmex* as an outgroup. While our *COI* tree provides a good framework to understand how microbes vary across groups of related ants, the exact branching patterns do not conform to the exact histories within this group.

### Microbial community analysis

For 86% of the processed ant workers (424 / 492), PCR reactions with eubacterial 16S rRNA primers resulted in the amounts of amplicon visibly exceeding those for negative controls. However, we noted clear differences among Dorylinae species in the proportion of workers for which 16S rRNA gene amplification succeeded. For example, we successfully amplified bacterial DNA from 98% of *Eciton burchellii* specimens (120 / 122), 90% of *Labidus praedator* specimens (69 / 77), but only 15% of *Labidus coecus* specimens (4 / 26) (Fig. 1). For three out of these four *L. coecus* workers, 16S rRNA amplicon sequencing resulted in datasets where a large majority of reads represented unique genotypes and OTUs that were also abundant in negative controls. This indicated that the numbers of bacteria in tissues of those ants were low, relative to the amounts of bacterial DNA present in reagents and laboratory environments (Salter *et al.* 2014). In contrast, 55 / 57 of *L. praedator* amplicon libraries, 78 / 79 *E. burchellii* libraries, 55 / 55 libraries prepared for other New World Dorylinae species and 13 / 15 for Old World Dorylinae were enriched in bacterial taxa rare in negative controls; furthermore, the majority of bacteria were previously known to colonize ants (Fig. S1). These findings suggest that army ant species differ in the numbers of bacterial cells they contain, although follow-up experiments using more accurate quantitative methods are required to confirm this observation.

We removed from analysis the eight libraries where the majority of reads represented genotypes and OTUs abundant in negative controls, and focused on the remaining 202 libraries (Fig. S1). After contaminant filtering and removal of unaligned, chimaeric or non-bacterial sequences, a total of 3,516,686 reads remained, with between 1,399-66,405 (median 15,466) reads per experimental library (Table S6). All calculations below are based on proportions of OTUs, as well as proportions of unique genotypes within OTUs, in individual libraries after the aforementioned quality control steps.

The species-level diversity of army ant-associated bacteria was low. Across all experimental libraries, the median relative abundance of the dominant 97% OTU was 93.9%, the median number of OTUs that reached at least 1% relative abundance in a given library was only two, and only 11% of specimens harbored more than three OTUs exceeding this abundance threshold (Tables S6–S8). Across all 202 libraries, six dominant OTUs accounted for an average of 92.6% reads. The two most abundant, representing the known specialized army ant symbionts Unclassified Firmicutes (OTU001) and Unclassified Entomoplasmatales (OTU002), accounted for an average of 57.8% and 21.6% of reads, respectively. However, there were consistent differences in microbial community composition across libraries representing different ant clades (Fig. 1). In three of the genera, *Eciton, Labidus* and *Nomamyrmex*, Unclassified Firmicutes and Unclassified Entomoplasmatales dominated microbial communities of a large majority of the sampled workers. Both of these microbes were also found in some, but not all, species of *Neivamyrmex* and Old World army ants. However, in many characterized ants other bacteria were quite common. For example, in some colonies of *Eciton* and *Labidus,* bacteria identified as Actinomycetales, Bartonellaceae, *Weissella* or *Spiroplasma* were abundant. Microbial communities of *Neivamyrmex* and *Aenictus* species were often dominated by *Wolbachia*, a widespread symbiont of insects that was strikingly absent from the other Dorylinae genera. Data for some species, such as *Eciton rapax* or *Neivamyrmex sumichrasti*, also raise the possibility for widespread domination of worker-associated microbial communities by a single specialized bacterial OTU.

In addition to variability across higher-level taxa (e.g. genera), we found intraspecific variation in microbiota across several scales within two well-sampled species, *Eciton burchellii* and *Labidus praedator* (Fig. 2). Permutational Analysis of Variance revealed that bacterial communities differed between these two species (ADONIS: F_1,123_ = 13.96, p = 0.001), but also between collection sites (F_7,116_ = 2.21, p = 0.005), though the interaction term was not significant (F_2,114_ = 2.45, p = 0.057; note that colonies of both species were only available from three locations). Furthermore, we found significant differences in microbial community composition among sympatric *E. burchellii* colonies in all cases in where at least three colonies were available from a single site (F_3,15_ = 3.38, p = 0.002 in Monteverde, Costa Rica; F_3,16_ = 4.78, p = 0.014 in Henri Pittier NP, Venezuela; and F_2,11_ = 3.29, p = 0.024 in Cocahoatan, Mexico). These differences were largely due to Unclassified Entomoplasmatales (OTU002), Actinomycetales (OTU005) and *Weissella* (OTU006) being abundant in some colonies of *E. burchellii* at the expense of Unclassified Firmicutes (OTU001) (Fig. 2, Table S9). We found no significant differences across colonies of *L. praedator* from either Monteverde (F_3,13_ = 1.83, p = 0.097) or from Henri Pittier NP (F_3,12_ = 0.89, p = 0.634).

**Figure 2.**
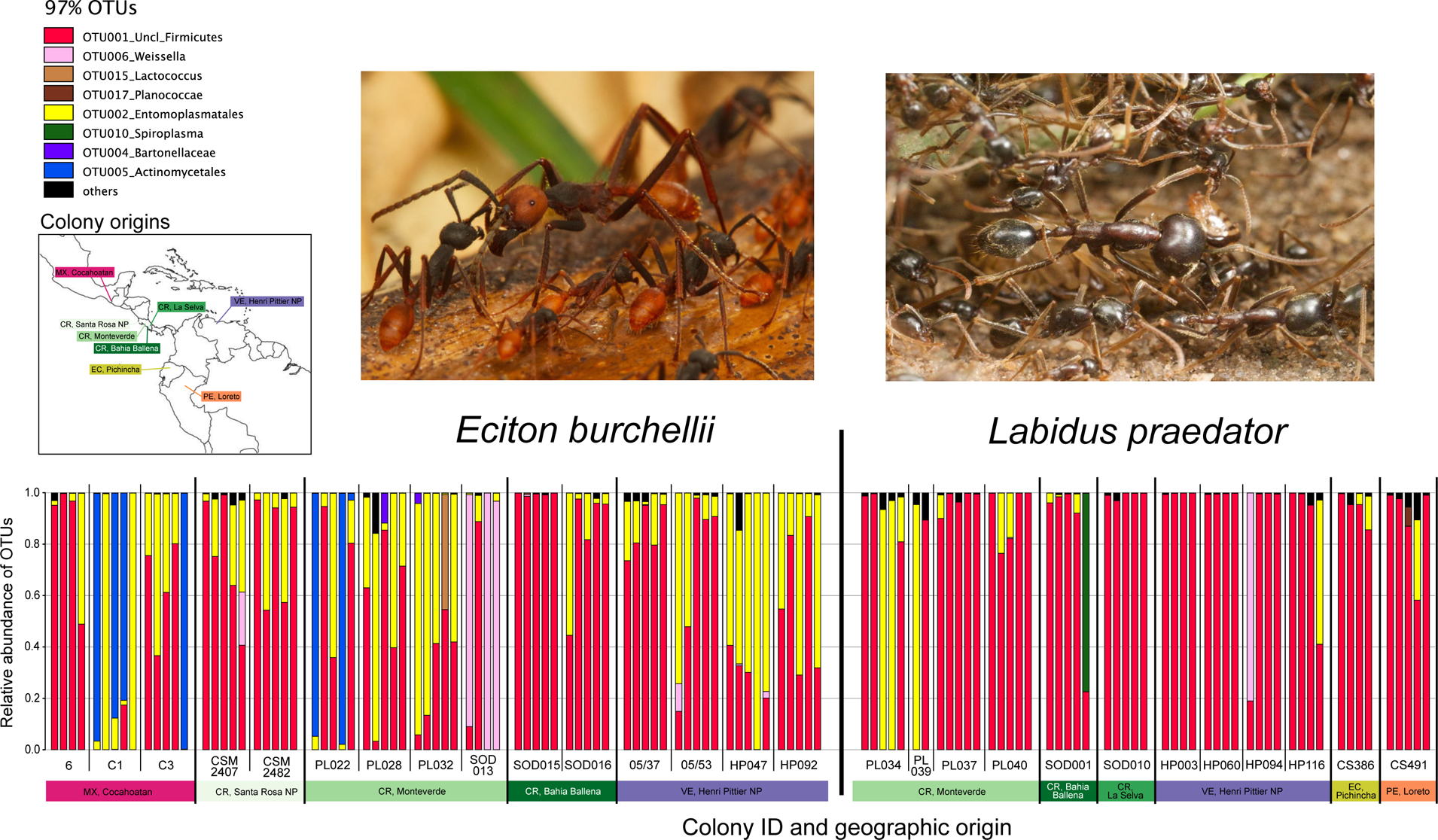
Variation in the microbial community composition at the colony and species level in two broadly sampled ant species. The relative abundance of microbial species (97% OTUs) is shown for individual workers from 15 colonies of *Eciton burchellii* and 12 colonies of *Labidus praedator*. Bars represent individual ants; colonies are separated by thin lines, and populations from different locations by thicker lines. Ant photographs were taken by JGS.

### Strain-level symbiont diversity and phylogenetics: Unclassified Firmicutes

We also looked at unique 16S rRNA genotype distributions within the dominant symbiont OTUs (Fig. S2). In the case of Unclassified Firmicutes, a single unique genotype typically made up 75-80% of that OTU; we believe that in most cases it represented the error-free sequence of the single bacterial strain present in that ant. The remaining 20-25% of the OTUs consisted of low-abundance genotypes, most of which differed at a single nucleotide position from the dominant genotype; we believe that the vast majority of these rare “unique genotypes” contained sequencing errors. However, in some cases two distinct unique genotypes were present at high abundance, suggesting the presence of more symbiont variants in a single ant, or perhaps differences between rRNA operons within the genome of a single strain. Several 16S rRNA genotypes from the Unclassified Firmicutes OTU were distributed across multiple host species and geographic locations. At the same time, some *Eciton burchellii* and *Labidus praedator* colonies harboured multiple 16S rRNA genotypes of these bacteria, with different putative strains typically distinguishing individual workers.

Sequencing the ribosomal protein gene *rplB* provided us with much higher strain-level resolution. We obtained clean and unambiguous sequence traces for approximately 90% of *rplB* sequences of Unclassified Firmicutes (253 sequences in total), and about 70% of Entomoplasmatales. Notably, we observed ambiguous peaks in *rplB* chromatograms in a large proportion of samples where 16S rRNA amplicon sequencing indicated the presence of more than one symbiont genotype. One of the challenges with inferences of multiple strain coinfection was suboptimal specificity of our primers, developed using genomic sequences of distantly related bacteria. This was evidenced by Entomoplasmatales primers yielding occasional amplification of *rplB* from other Mollicutes, and in a few cases, of Unclassified Firmicutes. As such, while multiple peaks in chromatograms may often imply co-existence of related strains within a single ant, this may not always have been the case.

A maximum likelihood phylogeny of Unclassified Firmicutes based on near-full length 16S rRNA sequences indicated that the symbionts of Dorylinae and two Ponerinae specimens represent a distinct and well-supported clade that belongs to a poorly characterized branch of Firmicutes (Fig. 3A). Analysis of the *rplB* gene enabled finer-scale comparisons of relationships among strains. Because of challenges in reconstructing the branching order for deeper parts of the tree, we constrained this phylogeny based on deep branch relationships in the 16S rRNA tree, keeping Ponerinae and most *Aenictus* and *Dorylus* symbionts outside of the main *Eciton-Labidus* symbiont clade (Fig. 3B, see also Fig. S4 for an unconstrained tree). In both the constrained and unconstrained versions of the *rplB* tree, symbionts originating from the same host species formed distinct and well-supported clades: three for *Eciton burchellii*, four for *Labidus praedator*, and typically more than one for less comprehensively sampled species. These species-specific clades commonly included strains from distant geographic locations. In many cases, replicate workers from a single colony hosted different symbiont genotypes, which belonged to different host-specific clades. For example, among eight workers from *Eciton burchellii* colony PL028 we identified four symbiont genotypes representing three divergent clades. Among six workers from *Labidus praedator* colony PL037 we found five genotypes representing three divergent clades (Fig 3B). Unfortunately, low resolution and poor support of deeper nodes in the tree limited our abilities to fully characterize relationships between these disparate lineages.

**Figure 3.**
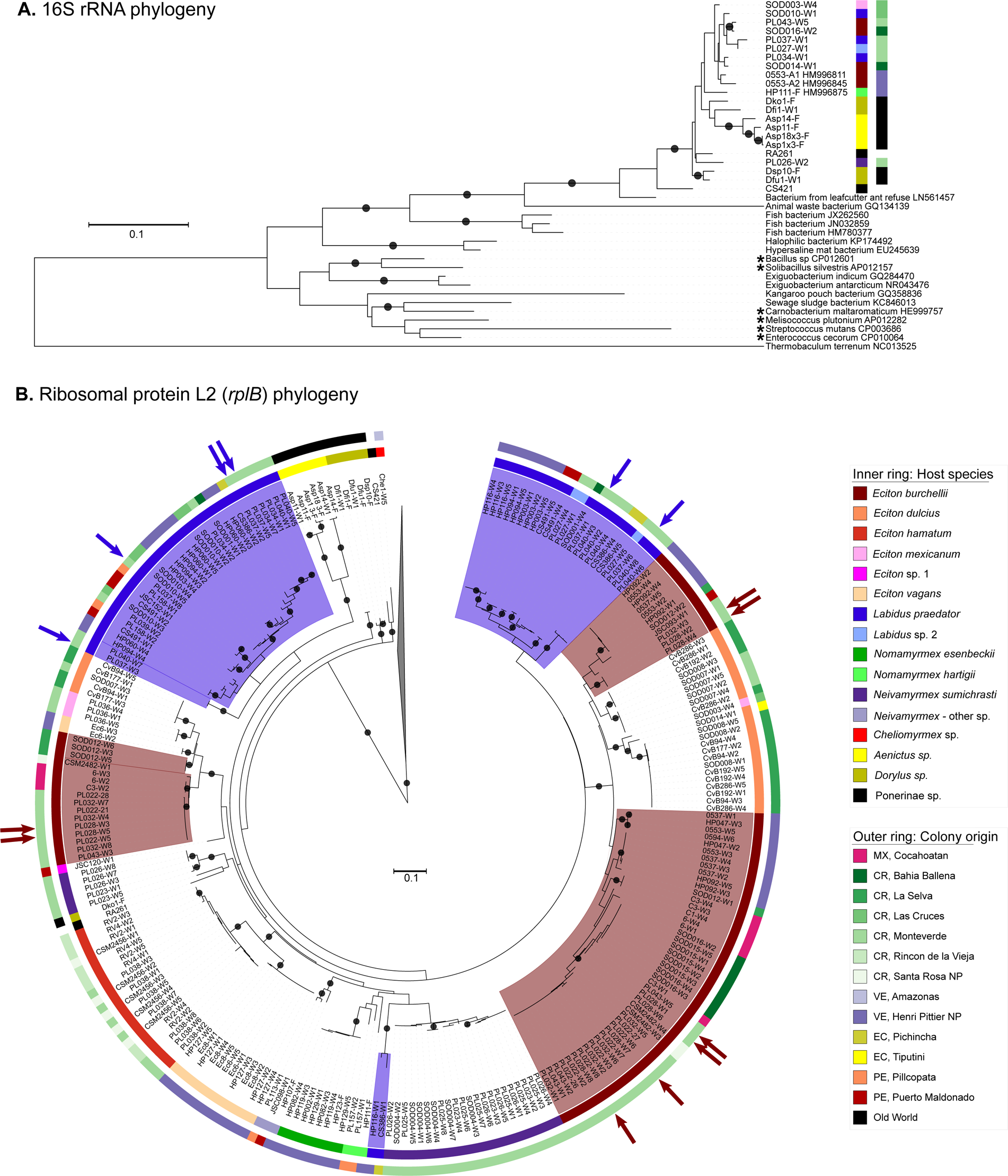
Phylogenetic position and relationships between Unclassified Firmicutes strains. A) ML phylogeny of selected Unclassified Firmicutes from diverse ants, as well as selected outgroups available in Genbank, based on near-full length sequence of the 16S rRNA gene. B) ML phylogeny of Unclassified Firmicutes from 252 ant workers, based on a 573bp sequence of ribosomal protein L2 (*rplB*) gene. Bootstrap support values of 95% or more are represented by black dots on branches. Inner colored ring indicates host species, and the outer ring indicates colony origin. In the *rplB* tree, we further highlighted clades composed of symbionts of the two most extensively sampled species, *Eciton burchellii* and *Labidus praedator*, and indicated with arrows all specimens from a single colony of each species. The six species indicated with asterisks in 16S rRNA tree were used as outgroups in *rplB* tree. Deeper nodes of *rplB* tree were constrained based on 16S rRNA tree, as described in Methods.

454 pyrosequencing of *rplB* amplicons provided a deeper insight into strain-level diversity of Unclassified Firmicutes in single hosts. Among the four sampled *E. burchellii* workers, all from a single colony, we obtained a total of 11,712 high-quality reads, with a range of 1360-5169 per sample. Sequences clustered into 43 OTUs at the 98% similarity level and each OTU was represented in only one worker. Within each library, between 99.54 and 99.58% of reads classified to a single OTU (Fig S3, Table S10), and in each of these four dominant OTUs, the consensus sequence was identical to the reference Sanger sequence from the same worker (Table S11). Analysis and manual inspection of alignments of reads classified to these four dominant OTUs strongly suggested that virtually all positions with under 99% consensus across reads were the result of sequencing errors at homopolymer sites, or of low sequence quality near the ends of reads (i.e. after the 300^th^ base; Table S11). Alignments of representative sequences from low-abundance OTUs suggested that sequencing error was the most likely driver of their existence: rare OTUs were unique to each sample and differed from sequences in common OTUs by multiple substitutions and indels, mostly in homopolymer tracks. Together, these results suggest that the four studied ants from a single colony hosted a single and unique strain of Unclassified Firmicutes each. Note that this may not always be the case, particularly in specimens where 16S rRNA amplicon sequencing data indicate the presence of different 16S rRNA genotypes in a single sample (Fig. S2) and where *rplB* traces obtained using the Sanger method are ambiguous.

### Strain-level symbiont diversity and phylogenetics—Unclassified Entomoplasmatales

In contrast to the patterns for Unclassified Firmicutes, our analyses of 16S rRNA amplicon sequence data suggested it was more common for single ants to harbour multiple genotypes of the Unclassified Entomoplasmatales symbiont (i.e. OTU002; Fig. S2). Noisy sequence traces from our efforts to sequence *rplB* gene fragments after amplification with Entomoplasmatales-specific primers were also consistent with multiple genotypes or strains in single workers. Phylogenetic placement of “clean” *rplB* sequences (Fig. 4) revealed that New World Dorylinae host multiple clades from this bacterial order, all quite distinct from named species in the genera *Entomoplasma, Mesoplasma, Mycoplasma* or *Spiroplasma*, for which genomic data are available. As seen in prior studies that relied on 16S rRNA (Anderson *et al.* 2012; Funaro *et al.* 2011), most army ant associates grouped into a single lineage, which in our analysis was made up exclusively of bacteria from these particular hosts. Within this lineage, symbionts typically grouped into host-specific clades. Well-sampled species such as *E. burchellii* harboured distinct symbiont genotypes, sometimes differing at a few nucleotide positions and other times belonging to divergent clades. But again, the fact that noisy sequence traces were commonly obtained when sequencing *rplB* product, combined with the fact that there was often more than one abundant 16S rRNA genotype within the Entomoplasmatales OTU (Fig. S2) in our amplicon dataset, suggests that individual workers may host multiple Entomoplasmatales strains relatively often.

**Figure 4.**
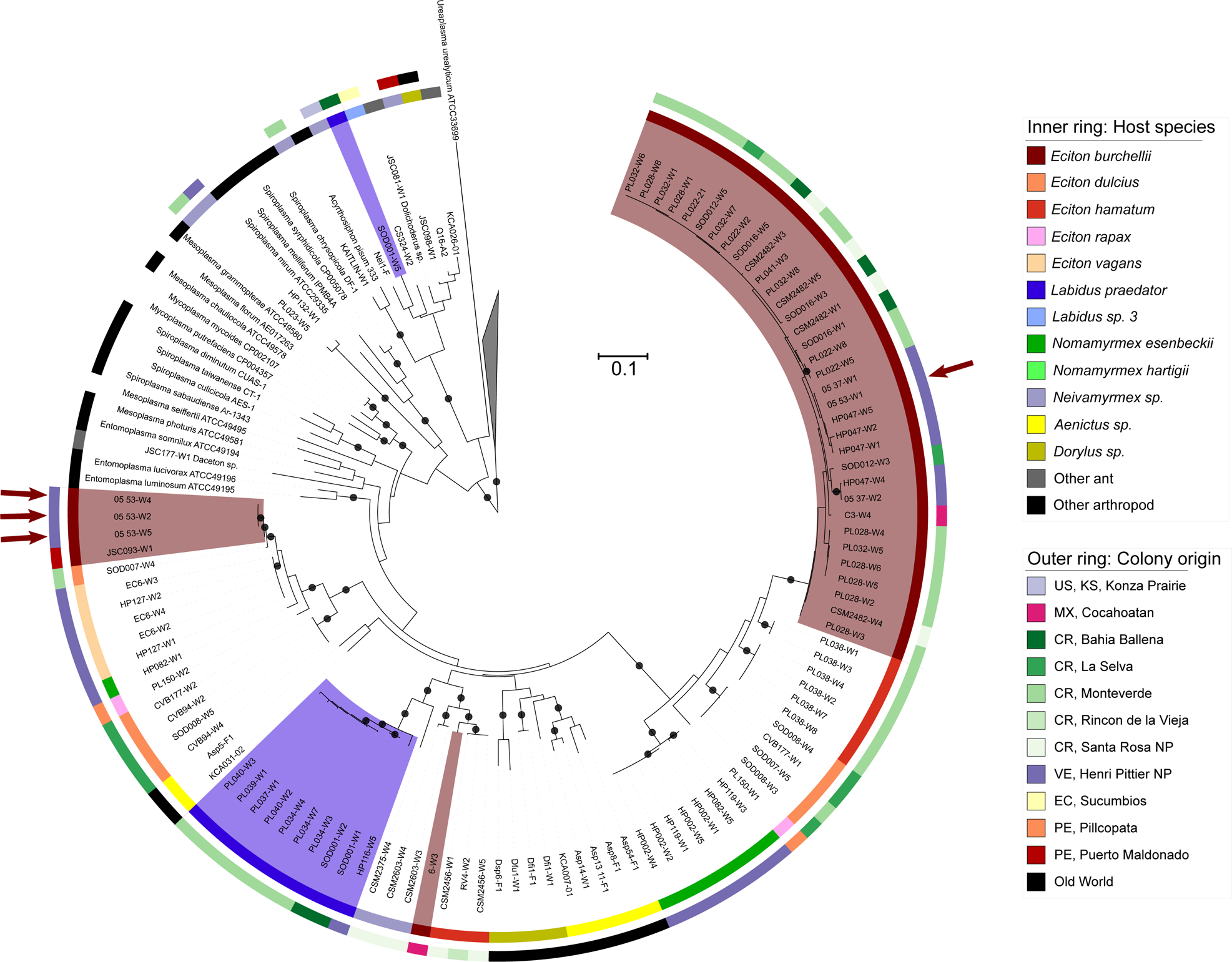
Diversity and specificity of Entomoplasmatales from Dorylinae and other ants, based on a 579bp sequence of ribosomal protein L2 (*rplB*) gene. Bootstrap support values of 95% and over are represented by black dots on branches of ML tree. Inner colored ring indicates host species and the outer ring - colony origin. Clades comprised of sequences amplified from the two most extensively sampled species, *Eciton burchellii* and *Labidus praedator*, are further highlighted. Arrows indicate all specimens from a single colony of *E. burchellii*.

### Fluorescence in situ hybridization

We observed clear and specific hybridization of fluorescently labeled probes (Fig. 5). Unlike in several ant species from other subfamilies (Sanders *et al.* in review), we observed bacteria colonizing the lumen of the pylorus and ileum in all examined Dorylinae species (Fig. 5. 1–3, 5.6). In *Eciton burchellii*, we identified several disjunct clusters of bacteria staining with either Entomoplasmatales or Unclassified Firmicutes probes (Fig. 5.3). In *Nomamyrmex hartigii*, most bacterial cells stained with Unclassified Firmicutes probes (Fig. 5.8), but we also found clusters staining with Entomoplasmatales probes (Fig. 5.5). In *Labidus praedator*, the signal of eubacterial probes overlapped perfectly with the signal of Unclassified Firmicutes probes, and we never detected Entomoplasmatales signal (Figs. 5.4, 5.7). Finally, in *Eciton rapax*, we observed no specific signal of Unclassified Firmicutes probes, and the signal of eubacterial probes overlapped with that of the Entomoplasmatales probe (Fig. 5.6). Interestingly, in this last species we found bacterial cells that stained with Entomoplasmatales probes in the foregut, within the crop, attached to what was likely a food pellet (Fig. 5.9). These cells formed dense clusters and were ovoid in shape, distinguishing them from rod-shaped Entomoplasmatales cells found in the hindgut.

**Figure 5.**
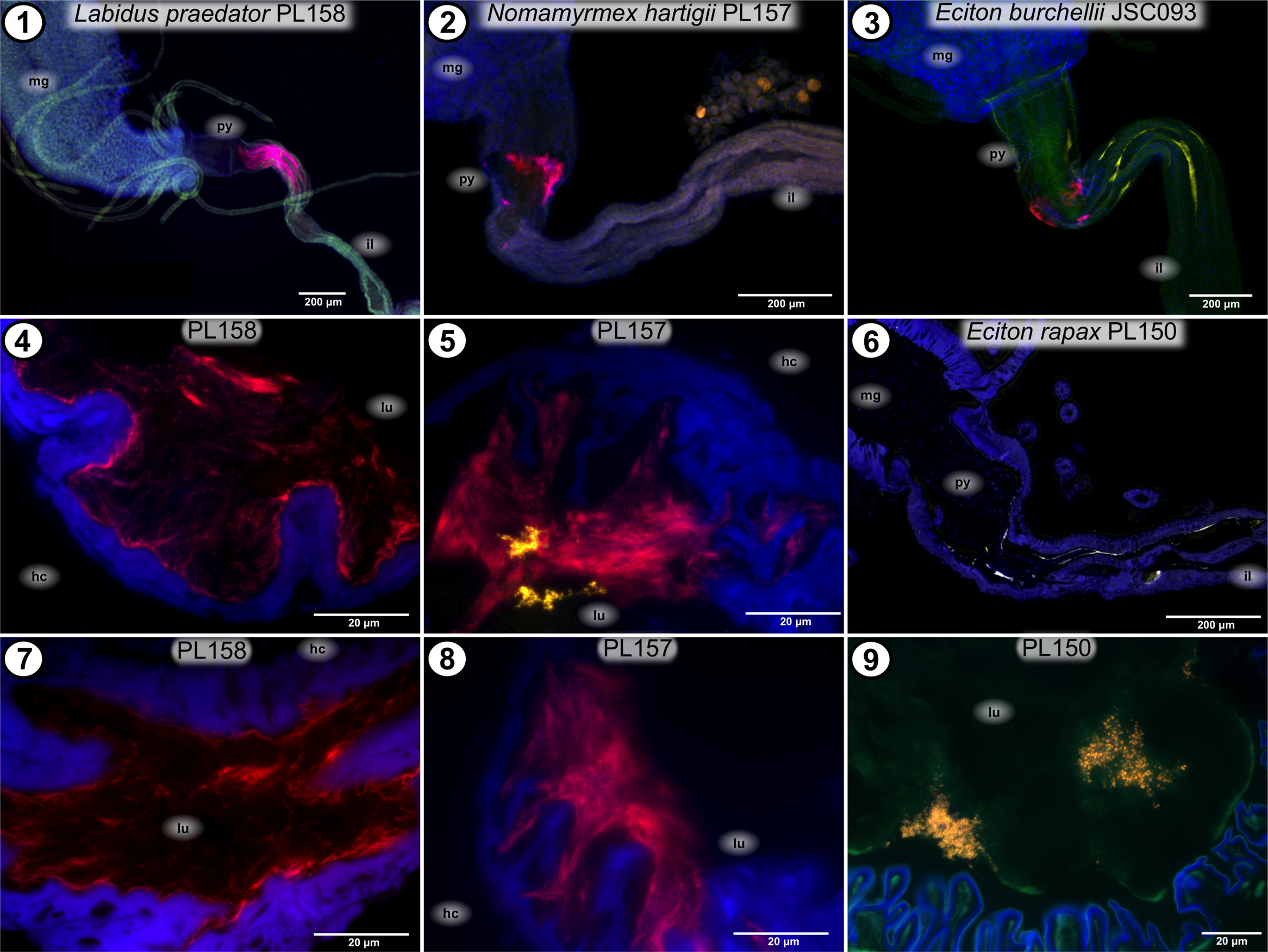
Localization of symbiotic bacteria within digestive tracts of workers of four species of Neotropical Dorylinae, demonstrated using fluorescent in situ hybridization (FISH), either of complete dissected guts (whole-mount – images 1-3) or of resin sections. Species and colony labels apply to all images below these labels. 1-3) Lower midgut and upper hindgut; 4-5) Cross-sections of the pylorus; 6) Longitudinal section of lower midgut and upper pylorus; 7-8) Section of ileum; 9) Bacterial colonies on a pellet in the foregut (crop). BLUE represents DAPI – universal DNA stain; GREEN – tissue autofluorescence (images 1-3 and 9 only); RED – either universal eubacterial probes (images 1, 2, 5) or Unclassified Firmicutes-specific probe (3, 4, 7, 8); YELLOW – Entomoplasmatales-specific probe (3, 5, 6, 9). Abbreviations: mg – midgut; py – pylorus; il – ileum; lu – gut lumen; hc – haemocoel.

The results for the four species are consistent with amplicon sequencing data, which revealed: a combination of Entomoplasmatales and Unclassified Firmicutes in most *E. burchellii* colonies, the dominance of Unclassified Firmicutes in guts of *Labidus praedator* and *Nomamyrmex hartigii*, and microbial communities dominated by distinct genotypes of Entomoplasmatales, without Unclassified Firmicutes, in single specimens from three *E. rapax* colonies (Fig. 1).

## Discussion

Our study combined broad sampling across populations of diverse army ants with deep sampling of two focal species. The specialized and broadly distributed microbes recovered through these efforts match those from prior studies relying on more traditional methods (Anderson *et al.* 2012; Funaro *et al.* 2011; Russell *et al.* 2012). However, the increased scale of sampling and the inclusion of a protein-coding gene in our phylogenetic analysis provide us with much greater insight into host specificity and evolutionary histories of the dominant symbiotic microbes. When combined with symbiont localization through fluorescence microscopy, our study identifies the two dominant, specialized bacteria as ectosymbionts, establishing army ants as yet another specialized feeding guild with conserved ectosymbioses.

### Army ant workers harbour low-diversity symbiont communities

In depth sequencing of 16S rRNA gene revealed that bacterial communities of the New World Dorylinae are dominated by a small number of specialized gut bacteria. While the gut microbiota of arthropods are less complex than those of vertebrates (Engel & Moran 2013), the low levels of diversity reported here still stand out. In a meta-analysis using cloned 16S rRNA libraries sequenced at varying depth, Colman et al. (2012) reported that different insect taxa and functional groups harbour 11-103 97% OTUs on average. Studies from a sample of diverse insects from Hawaii suggested that the average number of 97% OTUs at relative abundance of ≥1% was 7.5 (Jones et al. 2013). Herbivorous ants from the genus *Cephalotes* host 15-20 ‘core’ 97% OTUs (Hu *et al.* 2014). The number of ‘core’ 97% OTUs in guts of honeybees and bumblebees is approximately eight (Koch & Schmid-Hempel 2011a; Kwong & Moran 2015). This is much more than a median of only two 97% OTUs that we observed in army ants. It should be noted that these comparisons may have been somewhat impacted by differences in data analysis methods (Kunin et al. 2010; Quince et al. 2009). For example, the custom contaminant filtering used in the current study would have excluded real microbial symbionts of army ants that never exceeded ~0.2% of the community in any sequence library. However, the relative importance of such rare microbes to the host would be questionable, and proving that they represent real ant associates rather than experimental artifacts would be a daunting task.

### How ancient and specialized are the symbiotic gut bacteria of army ants?

Our data suggest that the two symbionts of army ants, Unclassified Firmicutes and Unclassified Entomoplasmatales, have persisted and often dominated the gut microbial communities of their hosts across vast phylogenetic and geographic scales. The Old World and the New World army ants split approximately 87 million years ago (Brady *et al.* 2014), and it appears that their common ancestor already hosted these two microbes. Further testing of this ancient infection model would benefit from examination of Dorylinae genera more closely related to the New World army ants than are either *Aenictus* and *Dorylus*, the two genera traditionally lumped within the “Old World” grouping (Borowiec 2016).

Another challenge to the model of ancient infection and specialization comes from the findings that divergent strains of Unclassified Firmicutes have been detected in predatory ants from the subfamily Ponerinae (Russell *et al.* 2009b; Fig. 3A), and that a microbe with 16S rRNA sequence identity of ~94% to New World Dorylinae symbionts was found in leafcutter ant refuse (Fig. 3A). Similarly, the large majority of Entomoplasmatales strains infecting Dorylinae belonged to army ant-specific clades (Funaro *et al.* 2011); but more recent efforts have detected closely related microbes in two Ponerinae species (Kautz *et al.* 2013a). These data suggest a broader distribution of these poorly characterized microbes, and some possibility for their independent acquisitions by army ants in the New World vs. Old World. More extensive sampling of microbes infecting other ants and non-ant hosts will provide more information about the evolutionary histories of the two bacteria. However, the deep phylogenetic branching within these ectosymbiont lineages and their cosmopolitan distributions across predatory ants argue for ancient origins of these symbionts, followed by histories of horizontal transmission among related hosts.

In contrast to these specialized bacteria, *Wolbachia* – the most widely distributed endosymbiont of insects (Zug & Hammerstein 2012) and the third most abundant microbe in our dataset - was prevalent in two distantly related and geographically disparate army ant genera (*Aenictus* and *Neivamyrmex*). It was strikingly absent from other army ants in our study. These patterns suggest more recent associations between army ants and their current suite of *Wolbachia* associates. Prior findings for close relatedness between *Wolbachia* of these ants and those of other ants and insects, indeed, argue against strong host specialization (Russell *et al.* 2009a). *Wolbachia* is known to vary in infection prevalence across host clades (Russell 2012; Russell *et al.* 2012), and the reasons for this are generally not understood. However, in *Anopheles* mosquitoes, bacteria of the genus *Asaia*, found in guts and other tissues, impede *Wolbachia* transmission (Hughes *et al.* 2014), raising the possibility that army ant ectosymbionts could shape *Wolbachia* distributions.

Beyond these three microbes, the distributions of other, less common bacteria across the sampled colonies did not follow clear patterns (Fig. 1). For example, deep sampling from two army ant species suggested differing abundance of *Actinomycetales* and *Weissella* species across colonies from the same locales, and sibling workers from the same colonies showed variability in the relative abundance of these taxa (Fig. 2). It is unlikely that these two bacteria have engaged in long-term or highly specialized associations with army ants. For example, BLASTn searches using representatives of the single *Weissella* OTU found in New World and the Old World army ants revealed 98-100% identity at the V4 region of 16S rRNA to other bacteria from animals, food products and other habitats. Representative sequences from the Actinomycetales OTU were 3% divergent from those of the closest known relatives, reported from corals, sponges and other environments, primarily marine. While abundant in three *E. burchellii* colonies, this microbe was absent from other ant samples, again suggesting a recent origin for this association.

The broad distribution and high relative abundance of specialized gut symbionts that appear to have co-diversified with their hosts for tens of millions of years is not standard insects. Indeed, bacterial gut communities of insects can be highly labile and strongly influenced by diet (Engel & Moran 2013). However, a number of social insects exhibit similar patterns. Honeybees and bumblebees provide one example. These two groups diverged ~87 million years ago (Cardinal & Danforth 2011), similar to the divergence time between New World and Old World army ants. Like army ants, these bees harbour specialized symbiotic gut bacteria that are thought to have persisted since the time of their common ancestor. Together with additional bacteria, the two most dominant ectosymbionts - *Snodgrassella* and *Gilliamella* – form conserved and stereotyped microbial communities (Engel *et al.* 2012; Kwong & Moran 2015). Both *Snodgrassella* and *Gilliamella* seem to have co-diversified with their bee hosts, albeit with multiple cases of apparent host switches and occasional losses (Koch *et al.* 2013). Similarly, the microbial communities of *Cephalotes* turtle ants, a genus that originated approximately 46 million years ago, consist of approximately 20 bacterial OTUs that are mostly shared across host species within this genus (Hu *et al.* 2014; Sanders *et al.* 2014). When clustered at an appropriate depths (i.e. 99% identity at 16S rRNA gene sequence), the community-level dendrogram for these microbiota shows congruence with the host phylogeny, providing a signal consistent with co-diversification. The complex microbial communities of termites also show evidence of co-diversification, at least at shorter evolutionary timescales (Brune & Dietrich 2015; Dietrich *et al.* 2014; Mikaelyan *et al.* 2015).

These three social insect groups feed on recalcitrant and imbalanced plant- or fungus-based diets and rely on microbial associates for digestion, nutritional supplementation (Brune & Dietrich 2015; Engel *et al.* 2012; Hu *et al.* in review; Warnecke *et al.* 2007) and in at least some cases, for defense (Koch & Schmid-Hempel 2011b). Omnivorous and carnivorous ants tend to host lower numbers of bacteria in their guts (Sanders *et al.* in review), and in the few groups characterized, the microbial communities appear to consist of less specialized associates (Hu *et al.* accepted; Ishak *et al.* 2011). While the abundance and importance of bacterial symbionts may vary across army ants, as suggested by differences in our 16S rRNA amplification success for different species (Fig. 1; see also Sanders *et al.* in review), the recurrent and ancient symbioses of Dorylinae appear exceptional among predatory insects.

### Community structure emerges at the strain level

While our 16S rRNA amplicon sequence data suggest differences in microbiota composition among army ant genera, many of the differences seen at regional and colony level scales were subtle. In addition, bacterial communities of sibling workers were often quite similar, with exceptions being driven by more sporadically occurring microbes. However, our *rplB* sequence data revealed that a good deal of genetic variability was undetected within the V4 region of 16S rRNA gene (Fig. S5A). Similar patterns were recently reported from corbiculate bees (Powell *et al.* 2016). The presence of distinct and well-supported host species-specific clades (Fig. 3B) suggests specialization and stability of host-symbiont associations across evolutionary timescales. These clades commonly consisted of strains from geographically distant locations, revealing further structure in our dataset not recovered with 16S rRNA. Furthermore, for each of the comprehensively sampled army ant species except *E. hamatum*, we identified several distinct host-specific symbiont clades, suggesting long-term co-existence of multiple divergent symbiont lineages, extending back for numerous generations within a single matriline.

Among our most curious findings was the discovery that distinct Unclassified Firmicutes strains colonize different workers, and that the majority of workers host only a single strain of this bacterium. While an individual workers’ microbiota may not be highly diverse, an unexpected range of symbiont strain diversity is retained within each army ant colony. While another group of social insect, honeybees, shows similar patterns of multi-strain persistence within colonies, different strains commonly infect the same bee workers (Engel *et al.* 2014; Powell *et al.* 2016), suggesting a unique type of diversity structuring within the army ants.

Similar to Unclassified Firmicutes, Entomoplasmatales of army ants are represented at the colony level by multiple distinct strains that often monopolize worker individuals (Figs. 4, S2C). However, the number of distinct host-specific clades appears to be less than in the case of Unclassified Firmicutes, and co-existence of two or more strains within a single worker seems to be more common. Findings that strain identities (Figs. 3 & 4) and overall ectosymbiont composition (Fig. 1) can vary among sibling workers raised the possibility for correlations between these levels of community structure. However, there appear to be no strong relationships between the genotype of the dominant ectosymbiont strain(s) and the species compositions of individual workers (Fig. S6A & S6B), leaving the drivers of ectosymbiont variation unexplained.

### For socially transmitted symbionts, modes of host colony founding are likely drivers of diversity and community structure

Sociality provides a unique context for the transfer of ectosymbionts, enabling regular spread of microbes among members of the same colony. Young honeybee and bumblebee workers acquire their specialized gut microbiota through exposure to feces present in a nest (Powell *et al.* 2014). Similarly, oral-anal trophallaxis seems to explain the transfer of gut symbionts in several herbivorous ants including those in the genus *Cephalotes*, whose callow workers consume fecal fluids from older adults (Cook & Davidson 2006; Lanan *et al.* 2016; Wilson 1976). It is plausible that newly eclosed Dorylinae workers acquire their microbiota through similar mechanisms (i.e. coprophagy or trophallaxis). High microbial strain-level diversity within army ant colonies may, thus, be maintained through a combination of social transmission and (as discussed below) the large numbers of individuals that found new colonies.

Modes of colony founding vary across social insects (Cronin *et al.* 2013), ranging from independent founding by single queens, to colony fission in which colonies split, enabling newly produced queens to initiate their reigns with a substantial work force. Army ants as well as honeybees engage in colony fission (Cronin *et al.* 2013; Peeters & Ito 2001). In contrast, queens of *Cephalotes*, fungus growing ants and bumblebees found colonies independently. Strain diversity for the ectosymbionts of *Cephalotes* ants (gut symbionts) and attines (residents of laterocervical plates on the cuticle - Mueller 2012) in some ways fit the expectations resulting from a strong symbiont bottleneck through a single individual. In particular, colonies seem differentiated by the presence of distinct strains within the core symbiont taxa (Anderson *et al.* 2012; Hu *et al.* 2014). For army ants, colony-level structuring of strain diversity is less apparent, at least based on our current levels of sampling depth. The founding of colonies by such large numbers of transmitting individuals provides a plausible explanation for the persistence of such diversity within colonies and, likely, numerous generations of a single matriline lineage. Colony founding differences have also been invoked as the driver behind higher *Snodgrassella* strain diversity among honeybees vs. bumblebees (Ellegaard & Engel 2016; Powell *et al.* 2016), making a case that the numbers of founding individuals may correlate with the diversity of socially transmitted symbionts at the colony level.

#### Symbiont strain competition, and a model-based approach

Transmission bottlenecks may, indeed, be much less severe for socially transmitted symbionts in social insects that utilize colony fission. But while this could explain the persistence of diverse symbiont strains within a single social unit, it does not explain why Unclassified Firmicutes strains should show such rare co-occurrence within single workers. One plausible explanation is that the initial inoculum received by newly eclosed workers tends to come from a single sibling. Under such a scenario, single strain domination would not require strong within-host microbial interactions as a mechanism behind such structure, especially if host ants become less receptive to colonization by future strains due to changes in behaviour, morphology, or physiology (Lanan *et al.* 2016). In contrast to this explanation, priority effects could drive some of the patterns observed for the army ant system; indeed, if first colonizers gained a strong competitive advantage, such symbionts could come to dominate their host resources. Host genotype could play an additional role (Spor *et al.* 2011), as could the large differences in behavior and anatomy among castes. It is also unclear if symbionts could turn-over within individuals over time. Distinguishing among these possibilities will require additional research.

Of further consideration will be theoretical efforts to understand the maintenance of symbiont diversity. For instance, strain distributions and persistence across worker generations could be modelled using population genetics tools, where symbiont strains act like alleles in populations (Jaenike 2012), represented here by colonies. Over time, as bacterial strains present in a colony monopolize newly emerged workers and old workers die, we would expect shifts in relative abundance of strains, extinction of some, and emergence of new strains through mutation. Horizontal transfer of strains among colonies and perhaps species would, similarly, simulate migration. Finally, colony fission events would have an effect comparable to emergence of geographic barriers. We reason that the large colony size and a lack of transmission bottlenecks suggest that genetic drift-like processes are a weaker force in eroding symbiont diversity in army ant colonies. With occasional novel strain acquisition through horizontal transfer and new strains arising through mutations, such high diversity could be maintained. The role of natural selection acting at the individual-, colony-or other levels must also be considered. Yet our current lack of knowledge on symbiont function inhibits our abilities to derive strong predictions.

#### What functional role do army ant gut microbes play?

Due to their stability over long evolutionary time and presence across multiple species our data raise the possibility for an important functional role for the most abundant gut microbes of army ants. Army ants are predatory, with diets consisting primarily of other insects (Kronauer 2009). While researchers did not foresee specialized symbioses for predatory ants, nitrogen-recycling symbioses found among some herbivorous ants (Feldhaar *et al.* 2007), were largely anticipated due to their reliance on nitrogen-poor diets (Cook & Davidson 2006; Davidson *et al.* 2003; Russell *et al.* 2009b). Differences in metabolic signatures have been seen for the gut microbiota of herbivorous and carnivorous mammals (Muegge *et al.* 2011), raising some expectations for the functional differences between microbiomes of herbivorous versus predatory ants. Diets with high protein-to-carbohydrate ratios may be harmful to animals including ants (Dussutour & Simpson 2009), and specialized gut symbionts of insect-feeders might be expected to mitigate these challenges. Other carnivorous insects, such as carrion beetles (Kaltenpoth & Steiger 2014) harbor bacteria some of which may co-diversify with hosts, making it possible that specialized ectosymbioses could be regular features beyond the herbivores and fungus-growing insects (Cafaro *et al.* 2011; Hulcr *et al.* 2012).

In addition to nutritional considerations, research in other symbiont systems suggests possible functions for the specialized symbionts of army ants. Detoxification represents one possibility, as do contributions toward development, physiology, and immune system development (Broderick *et al.* 2014; Buchon *et al.* 2009; Shin *et al.* 2011). Protective roles could also be significant. Indeed, microbes present on external body surfaces or within the digestive tract of insects can defend against parasites and pathogens (Currie *et al.* 1999; Kaltenpoth & Engl 2014; Kaltenpoth *et al.* 2005; Koch & Schmid-Hempel 2011b). The specialized gut bacteria of bumblebees protect hosts against a trypanosomatid intestinal parasite, with strong specificity between symbiont and parasite genotypes (Koch & Schmid-Hempel 2011b, 2012). To our knowledge, there are no data on microbial parasites of army ants; nevertheless, it remains plausible that their bacterial symbionts could play defensive roles.

## Conclusions

Characterizing the diversity, distribution and functions of microbial symbionts across insects is crucial to understanding the history and present-day biology of the most diverse group of animals on the planet. Ants, in particular, represent one of the most successful groups of insects in terms of their diversity and biomass. Microbial symbionts are thought to have facilitated their dominance of prey-poor rainforest canopies (Davidson *et al.* 2003). Additionally, symbiotic microbes have been clearly important in the fungus-farming ants, which are among the most influential insects of the tropics (Hölldobler & Wilson 2011). Our data suggest that ants from a third specialized dietary niche – top predators - are also enriched for highly conserved and specialized symbionts, raising important questions for the roles and impacts of microbial symbionts across insect predators and other carnivorous animals.

## Acknowledgements

We thank Christoph von Beeren and Kaitlin Baudier for providing ant specimens. This project was supported by NSF grants 1050360 to JAR and 1050243 to CSM, as well as JSPS Postdoctoral Fellowship (Short-term) PE13061 to PŁ.

## Data accessibility

Sanger sequences were deposited in Genbank (accession numbers KX982883-KX983349). Amplicon sequencing data were deposited in NCBI Short Read Archive (BioProject: PRJNA341802; 16S rRNA data: SRR4409391-4409626; Unclassified Firmicutes rplB data: SRR4343839-4343842). Colony and worker details are provided in the Supplementary Tables S1–S3. In Tables S1–S3, we also listed the above accession numbers by colony, worker individual and data type.

## Author contributions

PŁ and JAR designed the research. PŁ, JGS, CSM, DJK, SOD and JAR provided specimens. PŁ, JAN and YH generated molecular data. PŁ, with contributions from JGS, YH, CSM and JAR, analyzed the molecular data and illustrated the results. PŁ and RK performed fluorescence microscopy. PŁ and JAR wrote the manuscript, and all authors contributed to revisions.

## Supporting Information

**Appendix S1**. Supplementary information text with expanded supplementary table legends (Table S1–S11) and figure legends (Fig. S1–S6), details of amplicon sequencing data analyses, and FISH protocols.

**Table S1**. Details of the New World army ant colonies used in this project.

**Table S2**. Details of Old World army ant and other ant colonies for which some data were generated

**Table S3**. Details of individual worker samples from the experimental New World army ant colonies

**Table S4**. PCR primers and reaction conditions used in this project.

**Table S5**. Fluorescent microscopy probes used in this project.

**Table S6**. Summary of 16S rRNA amplicon sequencing analysis: library classification, and read numbers at various stages of the analysis.

**Table S7**. 16S rRNA amplicon sequencing data: unique genotypes remaining after contaminant and quality filtering. For each genotype, we provided sequence, OTU-level classification, taxonomy assignment, and information on the number of times it occurred in each library.

**Table S8**. 16S rRNA amplicon sequencing data: 97% OTUs. For each OTU, we provided a representative sequence, taxonomy assignment, and information on the number of times it occurred in each library.

**Table S9**. The list of OTUs that differ significantly in relative abundance when comparing between *Eciton burchellii* and *Labidus praedator*, across locations that these species were collected from, or among replicate colonies of *E. burchellii* from the same location.

**Table S10**. 454 pyrosequencing of *rplB* amplicon of Unclassified Firmicutes: distribution of 98% OTUs across experimental libraries

**Table S11**. Summary of genotype-level diversity data for Unclassified Firmicutes in individual ant workers, based on 454 pyrosequencing of *rplB* amplicon.

**Figure S1**. The effects of removal of putative contaminants on microbial community composition across libraries.

**Figure S2**. Genotype-level diversity of specialized symbionts of army ants, based on amplicon sequencing of the V4 region of the 16S rRNA gene.

**Figure S3**. Strain-level diversity of Unclassified Firmicutes in four individual workers from *Eciton burchellii* colony PL028, based on 454 pyrosequencing of the *rplB* amplicon.

**Figure S4**. Unconstrained ML phylogeny of Unclassified Firmicutes based on ribosomal protein L2 (*rplB*) gene.

**Figure S5**. *rplB*-genotype level diversity within 16S rRNA genotypes of the two specialized symbionts of army ants.

**Figure S6**. The relationship between *rplB* genotypes of Unclassified Firmicutes and the microbial community composition in two army ant species.

## References

Andersen SB, Hansen LH, Sapountzis P, Sørensen SJ, Boomsma JJ (2013) Specificity and stability of the *Acromyrmex*– *Pseudonocardia* symbiosis. Molecular Ecology 22, 4307–4321.

Anderson KE, Russell JA, Moreau CS, et al. (2012) Highly similar microbial communities are shared among related and trophically similar ant species. Molecular Ecology 21, 2282–2296.

Baumann P (2005) Biology of bacteriocyte-associated endosymbionts of plant sap-sucking insects. Annual Review of Microbiology 59, 155–189.

Bennett GM, Moran NA (2015) Heritable symbiosis: The advantages and perils of an evolutionary rabbit hole. Proceedings of the National Academy of Sciences 112, 10169–10176.

Billen J, Buschinger A (2000) Morphology and ultrastructure of a specialized bacterial pouch in the digestive tract of Tetraponera ants (Formicidae, Pseudomyrmecinae). Arthropod Structure & Development 29, 259–266.

Borowiec M (2016) Generic revision of the ant subfamily Dorylinae (Hymenoptera, Formicidae). ZooKeys 608, 1–280.

Boswell GP, Britton NF, Franks NR (1998) Habitat fragmentation, percolation theory and the conservation of a keystone species. Proceedings of the Royal Society B: Biological Sciences 265, 1921–1925.

Brady S, Fisher B, Schultz T, Ward P (2014) The rise of army ants and their relatives: diversification of specialized predatory doryline ants. BMC Evolutionary Biology 14, 93.

Brady SG (2003) Evolution of the army ant syndrome: The origin and long-term evolutionary stasis of a complex of behavioral and reproductive adaptations. Proceedings of the National Academy of Sciences 100, 6575–6579.

Broderick NA, Buchon N, Lemaitre B (2014) Microbiota-induced changes in *Drosophila melanogaster* host gene expression and gut morphology. mBio 5.

Broderick NA, Raffa KF, Goodman RM, Handelsman J (2004) Census of the bacterial community of the gypsy moth larval midgut by using culturing and culture-independent methods. Applied and Environmental Microbiology 70, 293–300.

Brune A, Dietrich C (2015) The gut microbiota of termites: digesting the diversity in the light of ecology and evolution. Annual Review of Microbiology 69, 145–166.

Buchon N, Broderick NA, Chakrabarti S, Lemaitre B (2009) Invasive and indigenous microbiota impact intestinal stem cell activity through multiple pathways in *Drosophila*. Genes & Development 23, 2333–2344.

Cafaro MJ, Poulsen M, Little AEF, et al. (2011) Specificity in the symbiotic association between fungus-growing ants and protective *Pseudonocardia* bacteria. Proceedings of the Royal Society B: Biological Sciences 278, 1814–1822.

Caporaso JG, Kuczynski J, Stombaugh J, et al. (2010) QIIME allows analysis of high-throughput community sequencing data. Nature Methods 7, 335–336.

Cardinal S, Danforth BN (2011) The Antiquity and Evolutionary History of Social Behavior in Bees. PloS One 6, e21086.

Chandler JA, Morgan Lang J, Bhatnagar S, Eisen JA, Kopp A (2011) Bacterial communities of diverse *Drosophila* species: ecological context of a host–microbe model system. PLoS Genetics 7, e1002272.

Colman DR, Toolson EC, Takacs-Vesbach CD (2012) Do diet and taxonomy influence insect gut bacterial communities? Molecular Ecology 21, 5124–5137.

Cook S, Davidson D (2006) Nutritional and functional biology of exudate-feeding ants. Entomologia Experimentalis et Applicata 118, 1–10.

Corby-Harris V, Pontaroli AC, Shimkets LJ, et al. (2007) Geographical distribution and diversity of bacteria associated with natural populations of *Drosophila melanogaster*. Applied and Environmental Microbiology 73, 3470–3479.

Cronin AL, Molet M, Doums C, Monnin T, Peeters C (2013) Recurrent evolution of dependent colony foundation across eusocial insects. Annual Review of Entomology 58, 37–55.

Currie CR, Scott JA, Summerbell RC, Malloch D (1999) Fungus-growing ants use antibiotic-producing bacteria to control garden parasites. Nature 398, 701–704.

Darriba D, Taboada GL, Doallo R, Posada D (2012) jModelTest 2: more models, new heuristics and parallel computing. Nature Methods 9, 772–772.

Davidson DW, Cook SC, Snelling RR, Chua TH (2003) Explaining the abundance of ants in lowland tropical rainforest canopies. Science 300, 969–972.

Dietrich C, Köhler T, Brune A (2014) The cockroach origin of the termite gut microbiota: patterns in bacterial community structure reflect major evolutionary events. Applied and Environmental Microbiology.

Douglas AE (2015) Multiorganismal insects: diversity and function of resident microorganisms. Annual Review of Entomology 60, 17–34.

Duron O, Bouchon D, Boutin S, et al. (2008) The diversity of reproductive parasites among arthropods: *Wolbachia* do not walk alone. BMC Biology 6, 12.

Dussutour A, Simpson SJ (2009) Communal nutrition in ants. Current Biology 19, 740–744.

Ellegaard KM, Engel P (2016) Beyond 16S rRNA community profiling: intra-species diversity in the gut microbiota. Frontiers in Microbiology 7, 1475.

Engel P, Martinson VG, Moran NA (2012) Functional diversity within the simple gut microbiota of the honey bee. Proceedings of the National Academy of Sciences of the United States of America 109, 11002–11007.

Engel P, Moran NA (2013) The gut microbiota of insects – diversity in structure and function. FEMS Microbiology Reviews 37, 699–735.

Engel P, Stepanauskas R, Moran NA (2014) Hidden diversity in honey bee gut symbionts detected by single-cell genomics. PLoS Genetics 10, e1004596.

Feldhaar H (2011) Bacterial symbionts as mediators of ecologically important traits of insect hosts. Ecological Entomology 36, 533–543.

Feldhaar H, Straka J, Krischke M, et al. (2007) Nutritional upgrading for omnivorous carpenter ants by the endosymbiont Blochmannia. BMC Biology 5, 48.

Ferrari J, Vavre F (2011) Bacterial symbionts in insects or the story of communities affecting communities. Philosophical Transactions of the Royal Society of London, Series B: Biological Sciences 366, 1389–1400.

Fuchs BM, Glöckner FO, Wulf J, Amann R (2000) Unlabeled helper oligonucleotides increase the in situ accessibility to 16S rRNA of fluorescently labeled oligonucleotide probes. Applied and Environmental Microbiology 66, 3603–3607.

Fukatsu T, Nikoh N (2000) Endosymbiotic microbiota of the bamboo pseudococcid *Antonina crawii* (Insecta, Homoptera). Applied and Environmental Microbiology 66, 643–650.

Funaro CF, Kronauer DJC, Moreau CS, et al. (2011) Army ants harbor a host-specific clade of Entomoplasmatales bacteria. Applied and Environmental Microbiology 77, 346–350.

Himler AG, Adachi-Hagimori T, Bergen JE, et al. (2011) Rapid spread of a bacterial symbiont in an invasive whitefly is driven by fitness benefits and female bias. Science 332, 254–256.

Holldobler B, Wilson EO (1990) The Ants. Belknap Press.

Hölldobler B, Wilson EO (2011) The leafcutter ants: civilization by instinct W. W. Norton & Company, New York, NY.

Hu Y, Holway DA, Łukasik P, et al. (accepted) By their own devices: invasive Argentine ants have shifted diet without clear aid from symbiotic microbes. Molecular Ecology.

Hu Y, Lukasik P, Moreau CS, Russell JA (2014) Correlates of gut community composition across an ant species (Cephalotes varians) elucidate causes and consequences of symbiotic variability. Molecular Ecology 23, 1284–1300.

Hu Y, Lukasik P, Sanders JG, et al. (in review) All hands on deck for nitrogen metabolism in an ancient, multi-partite symbiosis between turtle ants and gut bacteria. Proceedings of the National Academy of Sciences.

Hughes GL, Dodson BL, Johnson RM, et al. (2014) Native microbiome impedes vertical transmission of Wolbachia in Anopheles mosquitoes. Proceedings of the National Academy of Sciences 111, 12498–12503.

Hulcr J, Rountree NR, Diamond SE, et al. (2012) Mycangia of ambrosia beetles host communities of bacteria. Microbial Ecology 64, 784–793.

Ishak HD, Plowes R, Sen R, et al. (2011) Bacterial diversity in *Solenopsis invicta* and *Solenopsis geminata* ant colonies characterized by 16S amplicon 454 pyrosequencing. Microbial Ecology 61, 821–831.

Jaenike J (2012) Population genetics of beneficial heritable symbionts. Trends in Ecology & Evolution 27, 226–232.

Jaenike J, Unckless R, Cockburn SN, Boelio LM, Perlman SJ (2010) Adaptation via symbiosis: recent spread of a *Drosophila* defensive symbiont. Science 329, 212–215.

Jones RT, Sanchez LG, Fierer N (2013) A cross-taxon analysis of insect-associated bacterial diversity. PloS One 8, e61218.

Kaltenpoth M, Engl T (2014) Defensive microbial symbionts in Hymenoptera. Functional Ecology 28, 315–327.

Kaltenpoth M, Gottler W, Herzner G, Strohm E (2005) Symbiotic bacteria protect wasp larvae from fungal infestation. Current Biology 15, 475–479.

Kaltenpoth M, Steiger S (2014) Unearthing carrion beetles’ microbiome: characterization of bacterial and fungal hindgut communities across the Silphidae. Molecular Ecology 23, 1251–1267.

Kaspari M, O’Donnell S (2003) High rates of army ant raids in the Neotropics and implications for ant colony and community structure. Evolutionary Ecology Research 5, 933–939.

Kautz S, Rubin BER, Moreau CS (2013a) Bacterial infections across the ants: frequency and prevalence of *Wolbachia, Spiroplasma*, and *Asaia*. Psyche 2013, 11.

Kautz S, Rubin BER, Russell JA, Moreau CS (2013b) Surveying the microbiome of ants: comparing 454 pyrosequencing with traditional methods to uncover bacterial diversity. Applied and Environmental Microbiology 79, 525–534.

Koch H, Abrol DP, Li J, Schmid-Hempel P (2013) Diversity and evolutionary patterns of bacterial gut associates of corbiculate bees. Molecular Ecology 22, 2028–2044.

Koch H, Schmid-Hempel P (2011a) Bacterial communities in central European bumblebees: low diversity and high specificity. Microbial Ecology 62, 121–133.

Koch H, Schmid-Hempel P (2011b) Socially transmitted gut microbiota protect bumble bees against an intestinal parasite. Proceedings of the National Academy of Sciences 108, 19288–19292.

Koch H, Schmid-Hempel P (2012) Gut microbiota instead of host genotype drive the specificity in the interaction of a natural host-parasite system. Ecology Letters 15, 1095–1103.

Kronauer DJC (2009) Recent advances in army ant biology (Hymenoptera: Formicidae). Myrmecological News 12, 51–65.

Kunin V, Engelbrektson A, Ochman H, Hugenholtz P (2010) Wrinkles in the rare biosphere: pyrosequencing errors can lead to artificial inflation of diversity estimates. Environmental Microbiology 12, 118–123.

Kwong WK, Moran NA (2015) Evolution of host specialization in gut microbes: the bee gut as a model. Gut Microbes 6, 214–220.

Lanan MC, Rodrigues PAP, Agellon A, Jansma P, Wheeler DE (2016) A bacterial filter protects and structures the gut microbiome of an insect. ISME J 10, 1866–1876.

Letunic I, Bork P (2016) Interactive tree of life (iTOL) v3: an online tool for the display and annotation of phylogenetic and other trees. Nucleic Acids Research doi: 10.1093/nar/gkw290.

Liberti J, Sapountzis P, Hansen LH, et al. (2015) Bacterial symbiont sharing in *Megalomyrmex* social parasites and their fungus-growing ant hosts. Molecular Ecology 24, 3151–3169.

Martinson VG, Moy J, Moran NA (2012) Establishment of characteristic gut bacteria during development of the honeybee worker. Applied and Environmental Microbiology 78, 2830–2840.

McFall-Ngai M, Hadfield MG, Bosch TCG, et al. (2013) Animals in a bacterial world, a new imperative for the life sciences. Proceedings of the National Academy of Sciences 110, 3229–3236.

Mikaelyan A, Dietrich C, Köhler T, et al. (2015) Diet is the primary determinant of bacterial community structure in the guts of higher termites. Molecular Ecology 24, 5284–5295.

Moran NA, McCutcheon JP, Nakabachi A (2008) Genomics and evolution of heritable bacterial symbionts. Annual Review of Genetics 42, 165–190.

Muegge BD, Kuczynski J, Knights D, et al. (2011) Diet drives convergence in gut microbiome functions across mammalian phylogeny and within humans. Science 332, 970–974.

Mueller UG (2012) Symbiont recruitment versus ant-symbiont co-evolution in the attine ant–microbe symbiosis. Current Opinion in Microbiology 15, 269–277.

Oksanen J, Kindt R, Legendre P, et al. (2007) The vegan package. Community ecology package 10.

Oliver KM, Degnan PH, Burke GR, Moran NA (2010) Facultative symbionts in aphids and the horizontal transfer of ecologically important traits. Annual Review of Entomology 55, 247–266.

Oliver KM, Moran NA, Hunter MS (2005) Variation in resistance to parasitism in aphids is due to symbionts not host genotype. Proceedings of the National Academy of Sciences of the United States of America 102, 12795–12800.

Peeters C, Ito F (2001) Colony dispersal and the evolution of queen morphology in social hymenoptera. Annual Review of Entomology 46, 601–630.

Powell E, Ratnayeke N, Moran NA (2016) Strain diversity and host specificity in a specialized gut symbiont of honeybees and bumblebees. Molecular Ecology 25, 4461–4471.

Powell JE, Martinson VG, Urban-Mead K, Moran NA (2014) Routes of acquisition of the gut microbiota of *Apis mellifera*. Applied and Environmental Microbiology doi: 10.1128/aem.01861-14.

Quince C, Lanzen A, Curtis TP, et al. (2009) Accurate determination of microbial diversity from 454 pyrosequencing data. Nature Methods 6, 639–641.

R Core Team (2015) R: A language and environment for statistical computing. R Foundation for Statistical Computing, Vienna, Austria.

Roche RK, Wheeler DE (1997) Morphological specializations of the digestive tract of *Zacryptocerus rohweri* (Hymenoptera: Formicidae). Journal of Morphology 234, 253–262.

Russell JA (2012) The ants (Hymenoptera: Formicidae) are unique and enigmatic hosts of prevalent Wolbachia (Alphaproteobacteria) symbionts. Myrmecological News 16, 7–23.

Russell JA, Funaro CF, Giraldo YM, et al. (2012) A veritable menagerie of heritable bacteria from ants, butterflies, and beyond: broad molecular surveys and a systematic review. PloS One 7, e51027.

Russell JA, Goldman-Huertas B, Moreau CS, et al. (2009a) Specialization and geographic isolation among *Wolbachia* symbionts from ants and lycaenid butterflies. Evolution 63, 624–640.

Russell JA, Moreau CS, Goldman-Huertas B, et al. (2009b) Bacterial gut symbionts are tightly linked with the evolution of herbivory in ants. Proceedings of the National Academy of Sciences 106, 21236–21241.

Russell JA, Sanders JG, Moreau CS (accepted) Hotspots for symbiosis: Function, evolution, and specificity of ant-microbe associations from trunk to tips of the ant phylogeny (Hymenoptera: Formicidae). Myrmecological News.

Salter SJ, Cox MJ, Turek EM, et al. (2014) Reagent and laboratory contamination can critically impact sequence-based microbiome analyses. BMC Biology 12, 1–12.

Sanders D, Kehoe R, van Veen FJF, et al. (2016) Defensive insect symbiont leads to cascading extinctions and community collapse. Ecology Letters 19, 789–799.

Sanders JG, Lukasik P, Frederickson ME, et al. (in review) Dramatic differences in gut bacterial densities help to explain the relationship between diet and habitat in rainforest ants. Molecular Ecology.

Sanders JG, Powell S, Kronauer DJC, et al. (2014) Stability and phylogenetic correlation in gut microbiota: lessons from ants and apes. Molecular Ecology 23, 1268–1283.

Schloss PD, Westcott SL, Ryabin T, et al. (2009) Introducing mothur: open-source, platform-independent, community-supported software for describing and comparing microbial communities. Applied and Environmental Microbiology 75, 7537–7541.

Shin SC, Kim S-H, You H, et al. (2011) Microbiome modulates host developmental and metabolic homeostasis via insulin signaling. Science 334, 670–674.

Spor A, Koren O, Ley R (2011) Unravelling the effects of the environment and host genotype on the gut microbiome. Nature Reviews: Microbiology 9, 279–290.

Stamatakis A (2014) RAxML version 8: A tool for phylogenetic analysis and post-analysis of large phylogenies. Bioinformatics 30, 1312–1313.

Stoll S, Gadau J, Gross R, Feldhaar H (2007) Bacterial microbiota associated with ants of the genus Tetraponera. Biological Journal of the Linnean Society 90, 399–412.

Sudakaran S, Salem H, Kost C, Kaltenpoth M (2012) Geographical and ecological stability of the symbiotic mid-gut microbiota in European firebugs, *Pyrrhocoris apterus* (Hemiptera, Pyrrhocoridae). Molecular Ecology 21, 6134–6151.

Warnecke F, Luginbuhl P, Ivanova N, et al. (2007) Metagenomic and functional analysis of hindgut microbiota of a wood-feeding higher termite. Nature 450, 560–565.

Wernegreen J, Kauppinen S, Brady S, Ward P (2009) One nutritional symbiosis begat another: Phylogenetic evidence that the ant tribe Camponotini acquired Blochmannia by tending sap-feeding insects. BMC Evolutionary Biology 9, 292.

Wilson EO (1976) A social ethogram of the neotropical arboreal ant Zacryptocerus varians (Fr. Smith). Animal Behaviour 24, 354–363.

Winston ME, Kronauer DJC, Moreau CS (2016) Early and dynamic colonization of Central America drives speciation in Neotropical army ants. Molecular Ecology doi: 10.1111/mec.13846.

Yun J-H, Roh SW, Whon TW, et al. (2014) Insect gut bacterial diversity determined by environmental habitat, diet, developmental stage, and phylogeny of host. Applied and Environmental Microbiology 80, 5254–5264.

Zug R, Hammerstein P (2012) Still a host of hosts for *Wolbachia*: analysis of recent data suggests that 40% of terrestrial arthropod species are infected. PloS One 7, e38544.

